# A computational framework for characterizing normative development of structural brain connectivity in the perinatal stage

**DOI:** 10.1101/2023.03.10.532142

**Authors:** Yihan Wu, Ali Gholipour, Lana Vasung, Davood Karimi

## Abstract

Quantitative assessment of the brain’s structural connectivity in the perinatal stage is useful for studying normal and abnormal neurodevelopment. However, estimation of the structural connectome from diffusion MRI data involves a series of complex and ill-posed computations. For the perinatal period, this analysis is further challenged by the rapid brain development and difficulties of imaging subjects at this stage. These factors, along with high inter-subject variability, have made it difficult to chart the normative development of the structural connectome. Hence, there is a lack of baseline trends in connectivity metrics that can be used as reliable references for assessing normal and abnormal brain development at this critical stage. In this paper we propose a computational framework, based on spatio-temporal atlases, for determining such baselines. We apply the framework on data from 169 subjects between 33 and 45 postmenstrual weeks. We show that this framework can unveil clear and strong trends in the development of structural connectivity in the perinatal stage. Some of our interesting findings include that connection weighting based on neurite density produces more consistent trends and that the trends in global efficiency, local efficiency, and characteristic path length are more consistent than in other metrics.

## I. Introduction

The human brain undergoes significant and rapid developments in perinatal period (the several weeks just before and after birth) [1], [2]. It can be argued that this stage is the most dynamic and the most critical period in brain development [3], [4]. During this period, processes such as neurogenesis, neural migration, synapse formation, and axonal growth work in synchrony to form the brain microstructure and to lay the foundations of structural and functional brain networks [5], [6]. Interruption of normal brain development in this period can result in lifelong neurodevelopmental and psychiatric disorders [2], [7]. Quantitative assessment of brain’s structural connectivity at this stage can enhance our understanding of the development of cognitive and behavioral capabilities and improve the diagnosis, management, and treatment of neurological disorders [8], [9].

Diffusion-weighted magnetic resonance imaging (dMRI) has played an increasingly prominent role in studying the development of brain micro-structure, macro-structure, and structural connectivity in the fetal and neonatal periods [10], [3]. Unlike histological studies that are invasive and very expensive, dMRI enables assessment of the entire brain in utero in 3D, and it allows studying larger populations at much lower cost. Quantification assessment of brain’s structural connectivity with dMRI is one of its most exciting applications [11], [8]. Building on a local estimation of fiber orientation distribution, tractography techniques are used to trace virtual streamlines connecting different brain regions. These streamlines, often weighted by some measure of tissue micro-structure integrity, determine the strength of connections between a predefined set of brain regions/nodes. This “connectome” can be regarded as a mathematical *graph* and graph-theoretic measures can be computed to characterize it quantitatively [12], [13]. Although dMRI-based brain connectivity analysis suffers from important challenges, constant technical advancements have improved its accuracy and reproducibility [14], [15]. Moreover, our understanding of the potentials and limitations of this method have greatly improved and we can more reliably interpret the quantitative results offered by this method [16], [17]. As a result, quantitative structural brain connectivity analysis with dMRI has been extensively used to study brain development, maturation, aging, and degeneration (e.g., [18], [19]).

The great majority of these technical and scientific developments have been focused on the brains of children, adolescents, and adults. Mainly due to the challenges of perinatal imaging and a lack of reliable quantitative analysis tools and resources, comparatively far less research has been devoted to studying the structural connectivity at this early stage. Furthermore, because of methodological variations in computing the structural connectome, inherent limitations of dMRI, and high inter-subject variability, it has been difficult to establish normative references for longitudinal and population studies. This represents a critical gap in knowledge as it is well known that adult-like topological structures and a highly structured brain connectome develop very early in life [20], [21]. Hence, there is an urgent need for methods and resources to enable accurate and reproducible quantitative assessment of structural brain connectivity in the perinatal stage. Such methods and tools can significantly enhance our understanding of brain development at this stage and enable us to probe the neurodevelopmental processes that shape the structure and function of the brain for the rest of life.

In this paper, we propose a new methodology for analyzing normal development of brain’s structural connectivity in the perinatal stage. Our approach is based on accurate spatial alignment and averaging of data from cohorts of subjects with the same age. To ensure accurate alignment of white mater structures across subjects, we will perform the registrations based on diffusion tensor and fiber orientation distribution. The proposed approach reduces the impacts of the inter-subject variability and low data quality, both of which can be substantial in this period. Hence, it enables highlighting the main developments in the structural connectivity that occur due to brain maturation. We expect that this approach should produce normative structural connectivity metrics that can be used as references for reliable assessment and comparison of normal and abnormal brain development at this critical stage. We apply the new method on a large cohort of subjects scanned between 33 and 45 postmenstrual weeks and analyze several important metrics of structural brain connectivity.

## II. Methods

### A. Data

We used the MRI data from the developing Human Connectome Project [22]. We considered postmenstrual ages (PMAs) between 33 and 45 weeks. For PMA of 35 weeks, for example, we used subjects scanned between 34.5 and 35.5 postmenstrual weeks. For PMAs around 38 weeks, dHCP contained many more subjects than needed for our analysis. Our recent work and works of other researchers have shown very little or no gain when more than 15 subjects are used in each age group [23], [24]. Therefore, we used at most 15 subjects for each PMA. For the earliest age of 33 weeks only seven subjects were available, but that was still sufficient for building detailed high-quality atlases. The dMRI scans for each subject included 20 non-diffusion-weighted measurements and 280 diffusion-weighted measurements at three b-values of 400 (n=64), 1000 (n=88), and 2600 (n=128).

### B. Computational pipeline

Figure 1 shows the data processing pipeline for computing population-averaged age-specific connectomes. The pipeline has two main branches. One branch uses fiber orientation distribution (FOD)-based registration to compute a tractogram for each age. The other branch uses diffusion tensor-based registration to compute maps of micro-structural biomarkers. The FOD-based alignment could have been used to also compute atlases of micro-structural biomarkers. However, we found that a diffusion tensor-based registration that accounted for local alignment of white matter tracts resulted in more accurate results. Different steps of the pipeline are described below. Note that this pipeline is applied separately for each age group to compute a structural connectome for each age between 33 and 45.

**Fig. 1.**
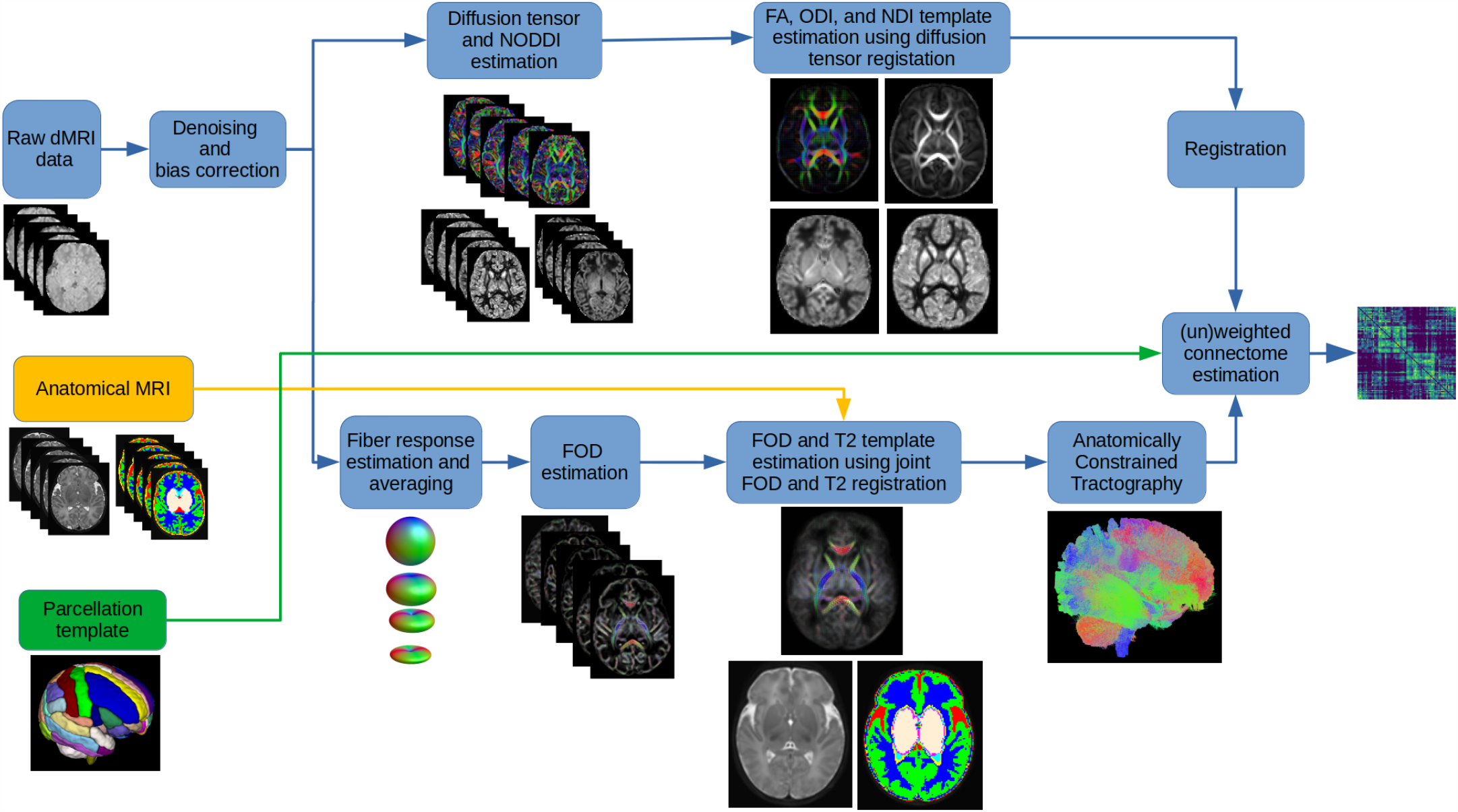
The proposed computational pipeline for computing age-specific structural connectomes.

#### 1) Data pre-processing

The dMRI data were first denoised using a PCA method [25], followed by B1 field inhomogeneity correction. Subsequently, all dMRI and anatomical data (i.e., T2 images and tissue segmentations) were resampled to an isotropic resolution of 1mm.

#### 2) Computing age-specific FOD templates and tractograms

We used the multi-tissue constrained spherical deconvolution method [26] for FOD estimation. This method is based on deconvolving the dMRI signal with signature response functions from white matter, gray matter, and cerebrospinal fluid. As suggested in [24], [27], we first estimated these response functions separately for each subject in an age group and then created an average response function. The average response function was used to estimate the FOD for each subject in the age group. A white matter FOD template was then estimated using symmetric diffeomorphic registration [28] of the white matter FOD maps of all subjects in the age group. The deformations computed based on the FODs were also used to deform the T2 images and tissue segmentation maps. Voxel-wise averaging and majority voting were used to estimate, respectively, a T2 template and a tissue segmentation template. Anatomically-constrained tractography [29] with a probabilistic streamline tracing method [30] was then applied using the FOD and tissue segmentation templates. Maximum angle between successive steps was set to 30 degrees and an FOD amplitude cut-off threshold of 0.01 was used as the stopping criterion. A total of five million valid streamlines were generated by seeding the entire brain.

#### 3) Computing age-specific templates of tissue micro-structure biomarkers

Proper weighting of edges in the structural connectome is an open problem. Although it is possible to compute edge weight/strength values based on tractography data alone, leveraging biomarkers of tissue micro-structure integrity is becoming more popular [15], [31]. In this work, we used biomarkers derived from the diffusion tensor and the Neurite Orientation Dispersion and Density Imaging (NODDI) models [32]. Specifically, for each subject in the age group we computed the fractional anisotropy (FA) using diffusion tensor fitting, and we computed the Orientation Dispersion Index (ODI) and the Neurite Density Index (NDI) from the NODDI model. We used the deep learning model proposed in [23] for computing the NODDI parameters. We then computed a template for these biomarkers using nonlinear diffusion tensor-based alignment [33]. These templates were then registered to the T2 template map for the same age group using affine registration. Note that the T2 and FOD templates were co-registered by design, as shown in Figure 1. Hence, after being registered to the T2 template, these biomarker templates could be used for weighting streamlines generated from the FOD template.

#### 4) Computing the structural connectome and connectivity metrics

To define connectome nodes, we used the Edinburgh Neonatal Atlas (ENA50) [34]. This atlas includes a parcellation of the neonatal brain gray matter into 100 regions. We registered this parcellation to our computed atas using deformable registration of the T2 image from the ENA50 atlas to the T2 atlas estimated by our pipeline for each age group. Using the gray matter parcellations as graph nodes and streamlines as edges, we computed structural connectomes. We used the SIFT2 algorithm to compute cross-sectional area multipliers to ensure the streamline densities reflect the density of the underlying white matter fibers. Additionally, we used the micro-structural biomarkers FA, NDI, and 1-ODI for connection weighting. The negative for ODI is standard practice and it is because ODI is a measure of fiber “dispersion”, whereas we would like to give a higher weight to higher microstructure “integrity”.

We computed six widely used structural connectivity measures: measures of network integration including characteristic path length (CPL) and global efficiency (GE), measures of network segregation including local efficiency (LE) and clustering coefficient (CC), small-worldness index (SW), and rich club coefficient (RC).

## III. Results and discussion

Figure 2 shows example FA and FOD atlases and tractograms. Overall, the results generated by our computational framework displayed displayed very high quality and detail. Table I shows the Spearman’s rank correlation coefficient of the six connectivity measures as functions of PMA, presented separately for the four different connection weighting schemes explored in this work. This coefficient quantifies the closeness of the relationship between two variables to a monotonic function; it is close to -1 or 1 if the relationship is highly monotonic.

**Fig. 2.**
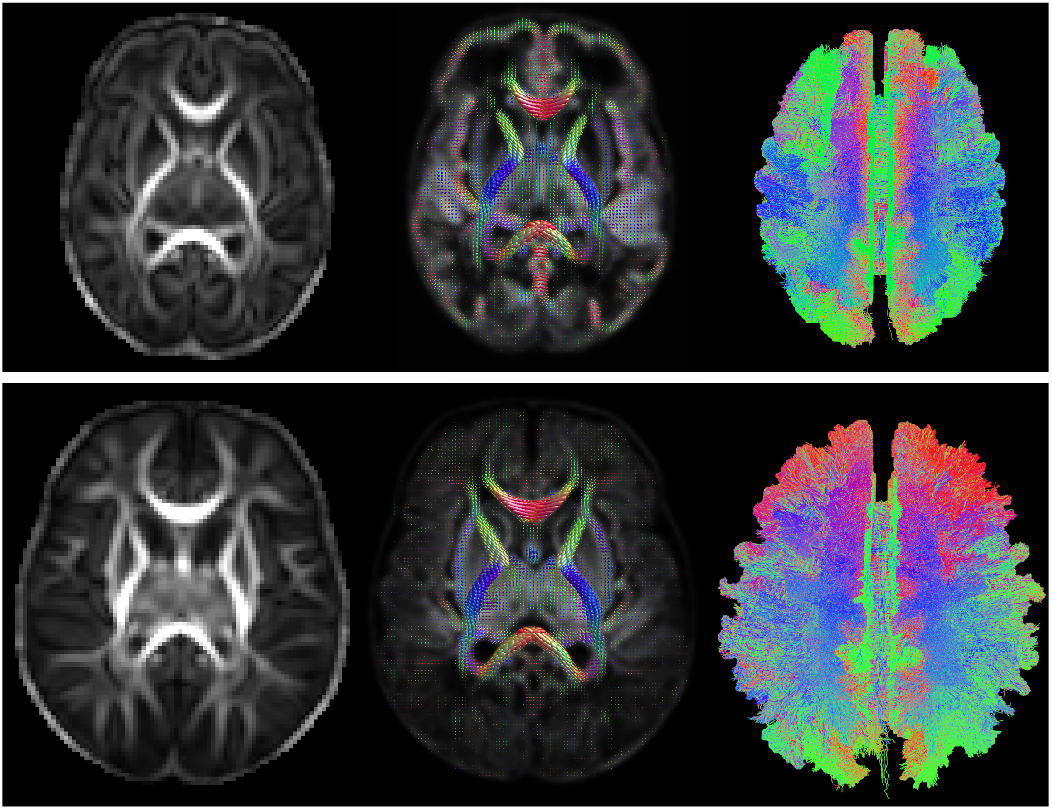
From left to right: example FA atlas, FOD atlas, and tractograms generated by our computational pipeline for 35 weeks (top) and 44 weeks (bottom).

**TABLE I.** Spearman’s rank correlation coefficient of different network connectivity measures as functions of PMA. Each row presents the coefficients for a different connection weighting scheme.

|  | GE | LE | CC | CPL | SW | RC |
| --- | --- | --- | --- | --- | --- | --- |
| FA | 0.99 | 0.97 | 0.09 | -0.99 | -0.64 | -0.15 |
| SIFT2 | 0.73 | 0.45 | -0.87 | -0.73 | -0.83 | 0.41 |
| 1-ODI | 0.19 | 0.10 | -0.01 | -0.21 | 0.36 | 0.16 |
| NDI | 0.98 | 0.97 | 0.97 | -0.96 | 0.29 | -0.92 |

An important and encouraging observation is the very strong relationship between the network connectivity measures and PMA, manifested by many Spearman coefficients that are very close to 1 or -1. The values shown in Table I are substantially higher than the values reported in prior studies that have analyzed the structural connectivity of individual subjects in this age range [35]. This shows the remarkable efficacy of the computational framework proposed in this paper to chart the normative development of structural brain connectivity in the perinatal stage.

Overall, the results show that GE, LE, and CPL have stronger monotonic relationships with PMA than the other three network measures. Furthermore, connection weighting based on NDI displays more consistent and stronger relationship with PMA. For all network measures considered here except SW, weighting of the connections based on NDI resulted in the highest (or very close to the highest) Spearman coefficient. Note that ODI and NDI are meant to provide more meaningful descriptors of the tissue micro-structure than FA, which is merely the degree of anisotropy of the best-fitted diffusion tensor. While ODI is related to the degree of dispersion of the neurites, NDI assesses the density of the neurites as the ratio of the diffusion signal contributed by the intraneurite compartment to that of the extra-neurite compartment. Although FA is a more commonly used measure, in part due to the lower data acquisition requirements of the diffusion tensor model compared with NODDI, our results show that weighting of the connections based on NDI produces the strongest and most consistent trends. Figure 3 shows the plots of the network measures as functions of PMA for the connectome that uses NDI-based edge weighting.

**Fig. 3.**
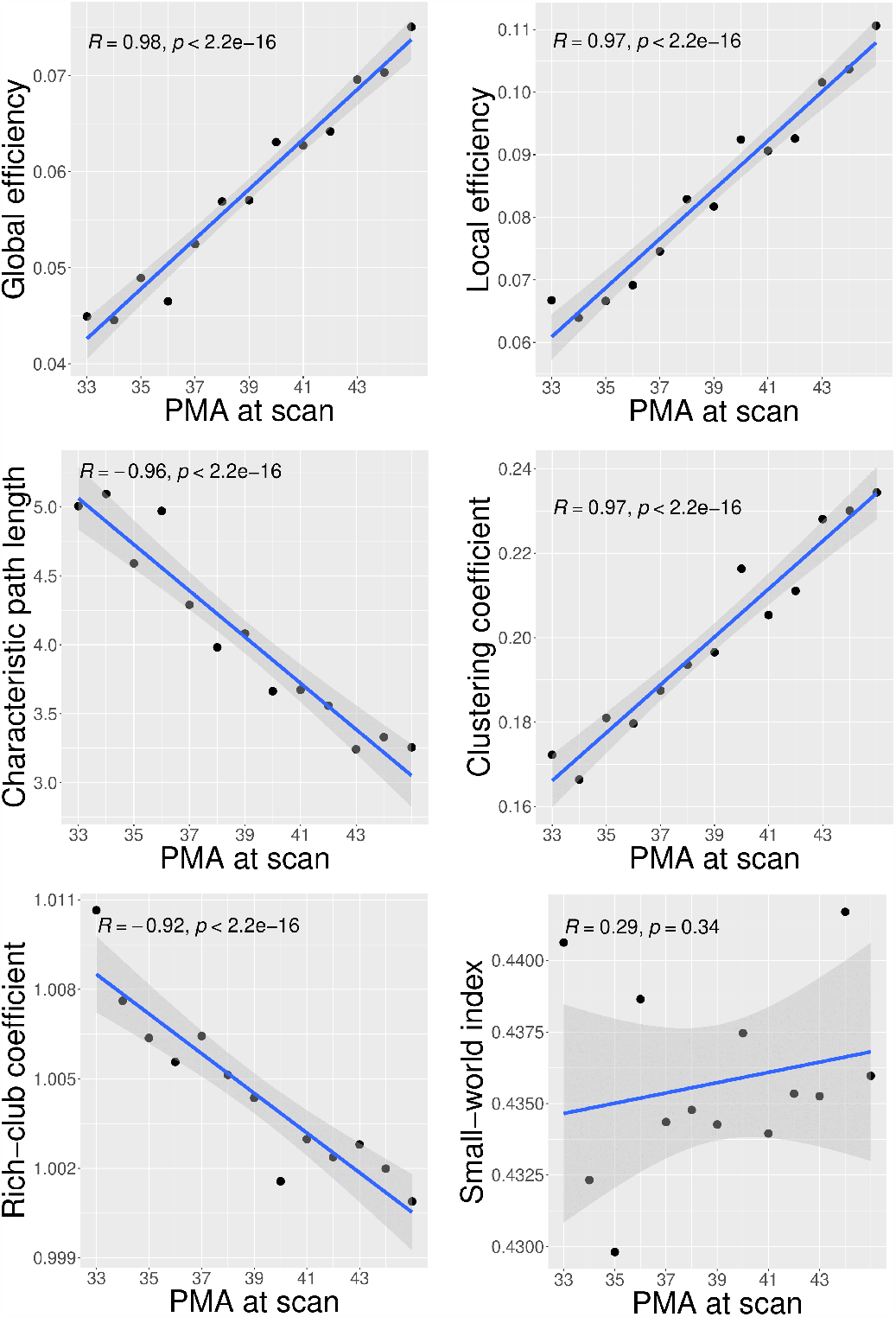
Plots of different structural connectivity measures versus PMA for the connectome edge weighting based on NDI.

Our observed trends in the structural connectivity metrics are interesting and conform with prior knowledge (e.g., [36], [37]). For example, they show a significant increase in network integration (in terms of GE and CPL), which indicates increased ability of the brain to integrate information from distant regions of the brain regions, and improved efficiency of communication between those regions. They also show a rapid increase in network segregation (in terms of LE and CC), which indicates the emergence of interconnected clusters of brain regions and increased ability of the brain to support information processing by such clusters.

## IV. Conclusions

This work has proposed a computational framework for studying the development of brain’s structural connectome in the perinatal stage. The new framework is based on accurate alignment of white matter structures across cohorts of subjects using tensor- and FOD-based registration. This enables reducing the inter-subject variability and reconstructing the developmental trajectories of the normal brain. Our results show that the proposed framework can unveil strong relationships between several critical measures of brain connectivity and PMA. Connectome edge weighting based on NDI was especially effective in uncovering strong and consistent trends in the structural connectivity measures. The developmental trends that have been reconstructed in this work can be used as reference baselines for comparing and contrasting normal and abnormal brain development in future works. Future works may also extend the proposed framework to analyzing brain connectivity in longitudinal and population studies.

## V. Acknowledgments

This study was supported in part by the National Institutes of Health (NIH) under grants R01EB031849, R01NS106030, and R01EB032366; and in part by the Office of the Director of the NIH under grant S10OD0250111.

The dHCP dataset is provided by the developing Human Connectome Project, KCL-Imperial-Oxford Consortium funded by the European Research Council under the European Union Seventh Framework Programme (FP/2007-2013)/ERC Grant Agreement no. [319456]. We thank the families who supported this trial.

## VI. Compliance with Ethical Standards

This research was conducted retrospectively using open access human data. MR images were acquired as a part of the dHCP which was approved by the National Research Ethics Committee and informed written consent given by the parents of all participants.

